# Transcriptome analysis of somatic cell populations in the *Drosophila* testis links metabolism and stemness

**DOI:** 10.1101/2020.02.24.962639

**Authors:** Silvana Hof-Michel, Christian Bökel

## Abstract

Due to its simple cellular architecture and genetic tractability the fly testis was one of the first model systems in which stem cell - niche interactions were studied at the molecular level. However, to date there is no comprehensive information on the endogenous, cell type specific transcription profiles associated with either stem cell or differentiated states. Focusing on the somatic lineage we have therefore isolated CySCs, differentiated CyCs, hub cells, and stem cell-like tumour cells overexpressing Zfh1, and have mapped their transcriptomes by RNAseq.

Here we report i) that the different somatic cell populations show extensive, genome wide differences in transcription levels, in particular associated with energy metabolism and innate immune signalling, ii) that differential activation of multiple signalling pathways renders Zfh1 overexpressing tumour cells unsuitable for use as a stem cell model, and iii) that the transcriptome data could be successfully used for identifying genes with stem cell specific expression patterns and for predicting aspects of stem cell physiology whose relevance for stem cell function could be validated in preliminary experiments.

The present data set should therefore facilitate future research on the interaction of stem cells with their niche using the highly successful fly testis model system.

## Introduction

At the tip of the fly testis, germline stem cells (GSCs) and somatic cyst stem cells (CySCs) surround a cluster of postmitotic somatic cells termed hub (Hardy et al., 1979) that provides the niche signals for the maintenance and activity of both stem cell populations (Fig. 1A). While the CySCs are the only proliferating somatic cell type in the adult testis (Amoyel et al., 2014), GSCs give rise to gonialblasts that undergo a further series of incomplete divisions resulting in 16-cell germline clusters that eventually enter meiosis and give rise to sperm.

**Figure 1:**
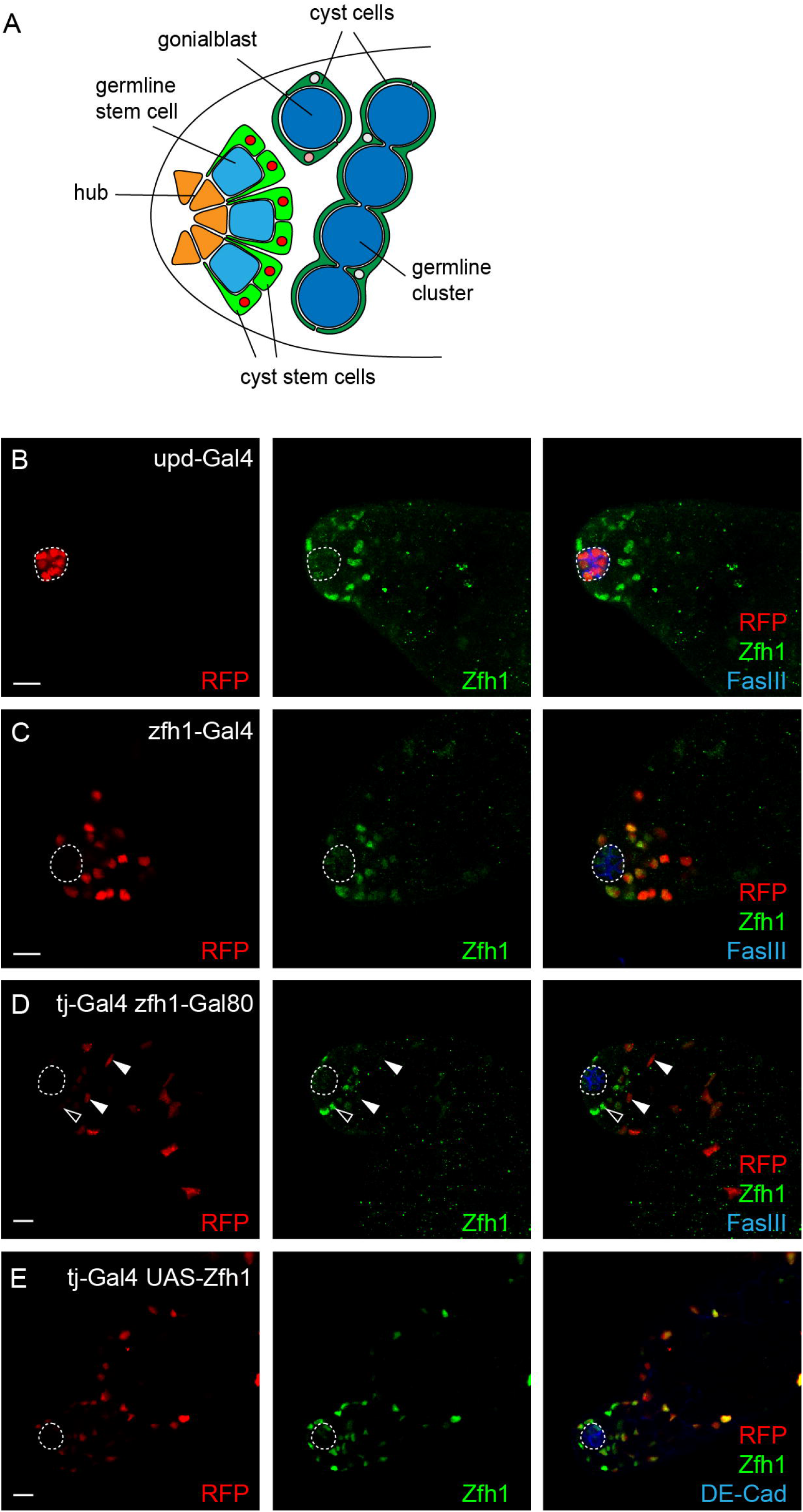
Labelling of somatic cell populations in the *Drosophila* testis. (A) Schematic representation of the testis tip. The hub (orange) is surrounded by germline stem cells (GSCs, light blue) and somatic cyst stem cells (CySCs, light green) that give rise to gonialblasts and differentiating germline clusters (dark blue) and their ensheathing cyst cells (CyCs, dark green). Zfh1 levels in the somatic lineage (nuclei marked red) drop sharply with differentiation. (B-E) Gal4-driven expression of a nuclear RFP (red) marks the hub (B, upd-Gal4), CySCs and their earliest CyC progeny (C, zfh1-Gal4), differentiated CyCs (D, tj-Gal4 zfh1-Gal80), and Zfh1-overexpression induced tumor-like cells (tj-Gal4 UAS-zfh1). Hub cells outlined by dashed line and marked in blue (FasIII), cyst stem cell nuclei in green (Zfh1). Hollow arrowheads in (D) denote Zfh1^high^ RFP^low^ and solid arrowheads heads Zfh1^low^ RFP^high^ cells. (B-E) Scale bars 10 μm.

Each testis contains about 10 to 14 GSCs and a similar number of CySCs (Amoyel et al., 2014; Hardy et al., 1979). The somatic CySCs already surround their germline counterparts and, following differentiation, two of their cyst cell (CyC) progeny ensheath each gonialblast. This close contact is retained as the germline clusters develop, and the CyCs eventually form a barrier between the germline clusters and the surrounding somatic tissues as they exit the niche (Fairchild et al., 2015; Fairchild et al., 2016). Division rates of the two stem cell types must therefore be coordinated to generate the appropriate number of differentiated cells (Amoyel et al., 2014). Despite the simple cellular organization this requires complex and bidirectional communication between the cells, making the *Drosophila* testis an ideal model system for studying stem cell niche interactions in a simple and genetically tractable model system (Losick et al., 2011).

However, the changes in expression level associated with either stem cell or differentiated status are only known for selected genes. Global and comprehensive transcriptome data are as yet available only at either cruder cellular resolution resolution, based e.g. on microarray analysis of RNA from microdissected testis tips (Cash and Andrews, 2012) or RNAseq of testes surgically split into three separate regions along their proximodistal axis (Vedelek et al., 2018). Clearly, RNA isolated in this way will come from a mixture of somatic and germline cells at various stages of differentiation.

Alternative approaches used microarrays (Terry et al., 2006) or RNASeq (Gan et al., 2010) on RNA extracted from whole testes in which particular, stem cell like cell populations were enriched through genetic manipulations. Similar approaches were also used in the ovary (Kai et al., 2005; Tiwari et al., 2019). This offers the advantage of genetic labelling and FACS separating germline and somatic before RNA isolation. However, even though the tumour-like populations generated by these genetic strategies express early markers suggesting an undifferentiated, stem cell like state, their relation to *bona fide*, endogenous stem cells remains unresolved (see also below).

Finally, genetically driven *in vivo* DamID (also called TaDa) (Southall et al., 2013) was applied to the fly testis to map PolII occupancy as a proxy for transcription in both the total somatic and germline lineages (Tamirisa et al., 2018). While this approach solves the problem of restricting the analysis to endogenous cells, PolII-DamID is notoriously less sensitive than methods relying on RNA isolation, which is reflected in the rather low number of genes scored as expressed. In addition, the Gal4 drivers chosen for this study did not allow differentiating between stem cells and differentiated cells in either lineage (Tamirisa et al., 2018).

To generate comprehensive, cell type specific transcriptome data sets for the somatic cells of the testis we have therefore genetically labeled various somatic cell populations in the testis, using specific combinations of Gal4 driver and Gal80 repressor lines to express a nuclear RFP. We were thus able to FACS isolate hub cells, CySCs, differentiated CyCs, and stem cell like cells overexpressing Zfh1 from live testes, and to map their transcriptomes by RNAseq.

Here we report that genes enriched in differentiated CyCs relative to somatic CySCs were found to be associated with mitochondrial biosynthesis and activity and an elevated oxidative energy metabolism. Consistent with preliminary observations of the mitochondrial oxidative state in the cyst cell lineage this difference may reflect a Warburg-type, glycolytically biased metabolic mode in the stem cells and, potentially, a metabolic division of labour between the differentiated CyCs and the germline clusters they ensheath.

In contrast, the set of genes differentially upregulated in stem cells was highly enriched for genes involved in innate immunity. In an accompanying, separate manuscript (Hof-Michel et al., 2020) we address the role of this immune signalling activity, and in particular the NF-kB transcription factor Rel, in stem cell competition.

Differential gene expression in hub cells relative to the adjacent CySCs, with which they share a developmental precursor again revealed expression differences in genes associated with glucose and energy metabolism.

Finallly, we compared the transcriptome of Zfh1 overexpressing cells with that of CySCs and their differentiated CyC progeny. Zfh1 overexpression induced similar and widespread changes, including the activation of multiple signalling pathways inactive in either endogenous population. Zfh1 overexpression also replicated only a minority of the differences between the endogenously Zfh1 positive stem cells and their Zfh1 negative progeny. The Zfh1 induced tumour cells thus appear to be of limited use as stem cell proxies.

We hope that the transcriptome data we present here will prove to be a useful resource for devising further, hypothesis driven research using the testis stem cell niche model system, in particular concerning the interplay between stemness, differentiation and metabolism.

## Results

### Labelling and isolation of somatic cell populations of the *Drosophila* testis for RNAseq

To follow up on our previous study of niche signal target genes in the somatic lineage of the *Drosophila* testis (Albert et al., 2018) we decided to isolate and purify various somatic cell populations and to compare their transcriptomes by RNAseq. We labelled the populations of interest by Gal4/Gal80^ts^ mediated expression of a nuclear DsRed protein termed RedStinger (Fig. 1B-E). Hub cells were marked by expressing RedStinger under upd-Gal4 control (Fig. 1B). The CySCs pool, presumably also including a fraction of their most recent, immediate progeny, was labelled using a Crispr/Cas9-generated zfh1-T2A-Gal4 genomic knockin line (Albert et al., 2018) (Fig. 1C). We used the Gal4-Gal80-Hack system developed by the Potter lab (Lin and Potter, 2016) to convert the zfh1-T2A-Gal4 driver into a zfh1-T2A-T2A-Gal80 repressor line (Fig. S1), which we combined with tj-Gal4 to label the entire CyC lineage with the exception of the Zfh1-positive stem cells (Fig. 1D). Finally, we used tj-Gal4 to co-overexpress Zfh1 with RedStinger, causing the Tj-positive somatic cells to proliferate (Fig. 1E). Note that under the 5d Zfh1 overexpression scheme of our experiment the proliferating somatic cells had not yet gained the ability to support germline stem cells, which, following even longer overexpression times, results in a mixed lineage, tumour-like phenotype (Leatherman and Dinardo, 2008).

We next hand dissected testes from flies containing the respective, RFP marked somatic cell populations. The labelled cells were then isolated by enzymatic digestion and mechanical disruption of the tissue, and five replicate samples per condition, each containing 125 to 150 fluorescent cells, were FACS sorted directly into RNAseq lysis buffer (Fig. S2). RNA was isolated from each sample, reverse transcribed, and PCR amplified before generating 20 separate cDNA libraries (4 tissue types x 5 replicas) for Illumina sequencing (for details, see methods section). Analysis of the raw sequencing data (trimming, quality control, mapping) was performed using the web-based A.I.R. RNAseq analysis package (Sequentia Biotech, Barcelona, Spain) (Vara et al., 2019). Sequencing yielded between 4.6×10^6^ and 9.1×10^6^ uniquely mapped reads per library (suppl. Table S1).

### Validation of the RNAseq results

Primary component analysis considering all expressed genes (suppl. Table S2) separated the twenty transcriptomes into four non-overlapping clusters corresponding exactly to the four starting cell populations (Fig. 2A). Spearman correlation between replicates of each of the four sample classes ranged from ρ=0.73 to ρ=0.96, demonstrating that the labelling and sorting of the cells had worked reproducibly (Fig. S3). Correlation between replicates was generally higher for samples where the starting material contained several hundred labeled cells per testis (differentiated CyCs and Zfh1-induced tumours) than for samples where a single testis contained only tens of labeled cells (hub and Zfh1 positive CySCs / early CyCs).

**Figure 2:**
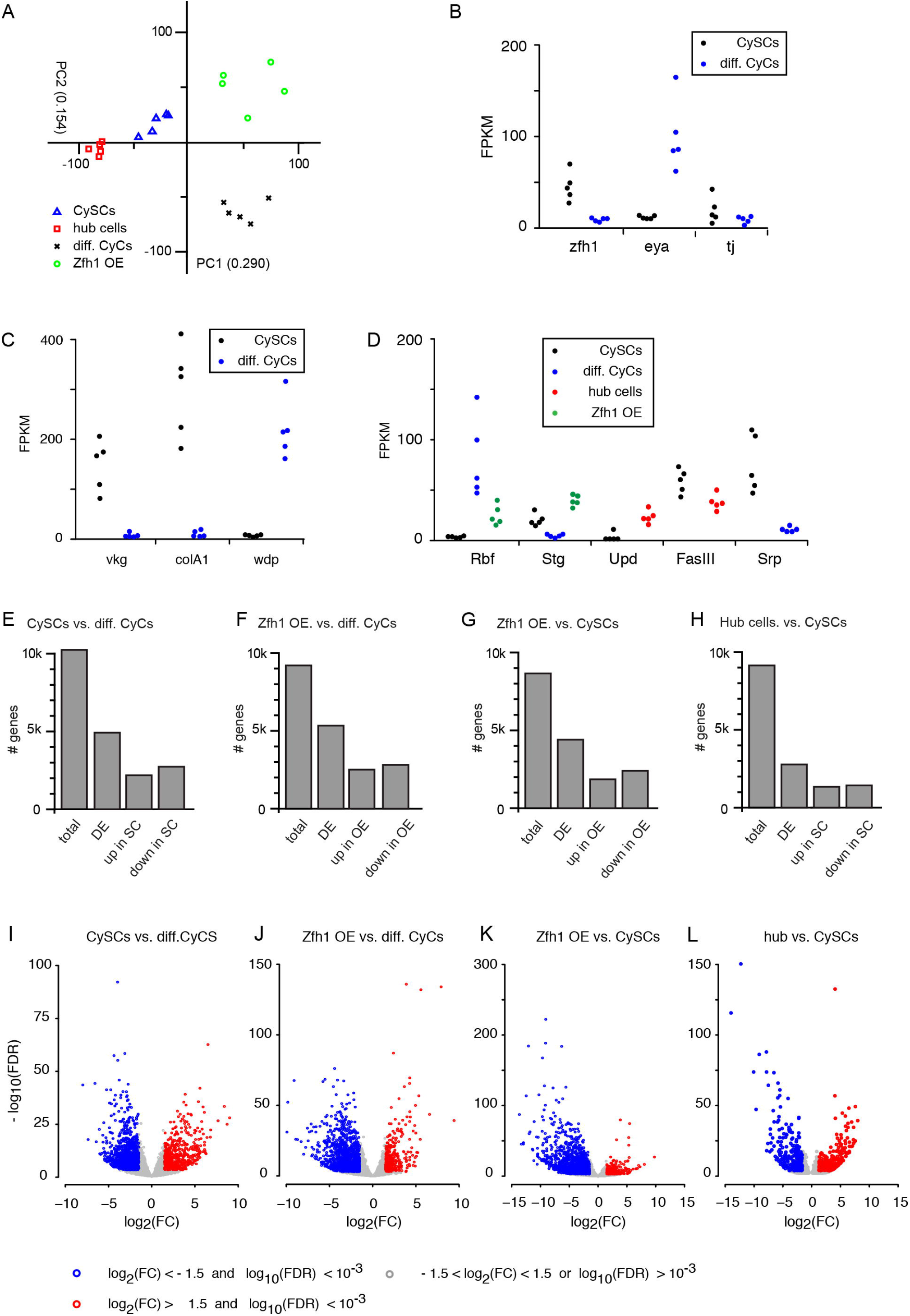
RNA-Seq of the somatic cell populations of the testis. (A) Primary component analysis of full transcriptomes for all 20 libraries. Each data point corresponds to one replica. Replicas from the four samples are cleanly separated into non-overlapping clusters. (B-D) Expression levels of candidate and marker genes in fragements per kilobase and 10^6^ reads (FPKM). Each data point corresponds to one replica. (E-H) Number of genes retained after low expression filtering (total) or scored as differentially expressed (DE), up- or downregulated according to DESeq2 with a false discovery rate (FDR) < 0.05 cutoff for comparisons of CySCs vs. differentiated CyCs (E), Zfh1 overexpressing cells vs. differentiated CyCs (F), Zfh1 overexpressing cells vs. CySCs (G), and hub cells vs. CySCs (H). (I-L) Volcano plots for the comparisons in (E-H) illustrating the more stringent cutoffs for fold change (FC) and FDR subsequently adopted. Downregulated (blue) and upregulated genes (red) are indicated.

A first pass inspection of the transcriptome data confirmed that expression levels of genes characteristic for the different cell populations were consistent with an overall successful separation. Endogenous Zfh1 transcript was detected at a 5.4-fold higher level in the stem cell samples relative to the differentiated CyCs (Fig. 2B). Consistent with previous observations (Li et al., 2003; Schulz et al., 2002), the Zfh1 positive CySCs also exhibited 2.2-fold higher expression levels of the general CyC lineage transcription factor Tj than their differentiated, Zfh1 negative CyC progeny (Fig. 2B). Similarly, mRNA levels of the Collagen IV subunit Vkg that, at the protein level largely restricted to the CySCs was increased 70-fold in the stem cells compared with their differentiated progeny (Fig. 2C). Conversely, expression of the differentiation marker Eya (Fabrizio et al., 2003) was upregulated 6.2-fold in differentiated CyCs relative to their stem cell progenitors (Fig. 2B).

In addition, we noticed several genes whose expression pattern in the testis was as yet undescribed, but whose molecular function was consistent with previously described niche and stem cell activities. For example, the collagen IV subunit Col4A1 was enriched 60-fold in the stem cells relative to the differentiated CyCs (Fig. 2C), mirroring the distribution of its binding partner Vkg. Conversely, in differentiated CyCs the transmembrane protein Wdp, a negative regulator of the Upd receptor dome (Ren et al., 2015), was expressed at roughly 20-fold higher levels compared to CySCs (Fig. 2C), conceivably preventing differentiated cells from responding to the Upd niche signal.

Consistent with CySCs being the only endogenously proliferating somatic cells in the testis, the cell cycle promoting kinase Stg was expressed at 1.6-fold higher levels in the stem cells (Fig. 2D), while expression of the anti-proliferative tumour suppressor Rbf was increased 12.5-fold in the non-dividing CyCs (Fig. 2D), confirming a previous report (Greenspan and Matunis, 2018). As expected from their induced proliferative state, Rbf was also downregulated and Stg expression increased in cells overexpressing Zfh1 (Fig. 2D).

In the hub, expression of the niche ligand Upd (Kiger et al., 2001; Tulina and Matunis, 2001) was increased 8.7-fold relative to somatic stem cells (Fig. 2D). Surprisingly, the homotypic adhesion molecule Faslll that tightly outlines the hub was expressed both in the hub and, more strongly, in the adjacent stem cells (Fig. 2D).

Finally, we already spotted multiple genes that possessed expression patterns suggestive of a role in stem cell regulation, but for which no function in the *Drosophila* testis niche had as yet been described. This class included e.g. the *Drosophila* GATA factor Srp (Fig. 2D) (see also below).

### Differential gene expression in the somatic lineage of the testis

Since the preliminary analyses described above suggested that our isolation protocol had successfully separated various somatic cell types of the testis, we proceeded with analyzing the overall gene expression profiles. Reads were analyzed with help of the A.I.R. package (Sequentia Biotech, Barcelona) (Vara et al., 2019). Low expression cutoffs were set by calculating global Jaccard indices for each comparison and discarding low expressed genes until correlation between replicates was maximized (Rau et al., 2013). Genes passing this filtering step were then tested for differential expression using the DESeq2 algorithm (Love et al., 2014), initially using a relaxed false discovery rate (FDR) of 0.05 as cutoff. Using these parameters, in each of the three pairwise comparisons between stem cells, differentiated cells, and Zfh1 overexpressing cells, around 10.000 of the 17.559 genes annotated in the BDGB Dmel6 genome release (dos Santos et al., 2015) were retained as expressed and, of these, between around 4.300 and 5.200 scored as differentially expressed (Fig. 2E-G). The degree of differential expression was somewhat lower (2.700 genes) for the comparison of CySCs and hub cells (Fig. 2H). The differentially expressed genes are listed in the supplementary information (suppl. Tables S3-S5). Heat mapping the zFPKM scores of these genes using an Euclidean distance metric for each comparison clustered the replicates of the same cell type, confirming reproducibility of the sorting protocol (Fig. S4). However, to increase reliability, we heuristically chose more stringent criteria, demanding both a DESeq2 FDR < 10^−3^ and an absolute of the log_2_(fold change) > 1.5 before scoring a gene as differentially expressed (Fig. 2I-L).

Using these stringent cutoffs, Zfh1 overexpression in the CyC lineage resulted in the upregulation of 362 genes and the downregulation of 1245 genes compared with the endogenously Zfh1 positive CySCs, and 349 upregulated and 1189 downregulated genes relative to the Zfh1 negative differentiated CyCs (suppl. Table S3). The two sets of differentially expressed genes showed significant overlap (p=0, hypergeometric test) (Fig. 3A). As already suggested by the PC analysis (Fig. 2A) these tumour-like cells thus form a unique population that is distinct from either cell type of the endogenous cyst cell lineage.

**Figure 3:**
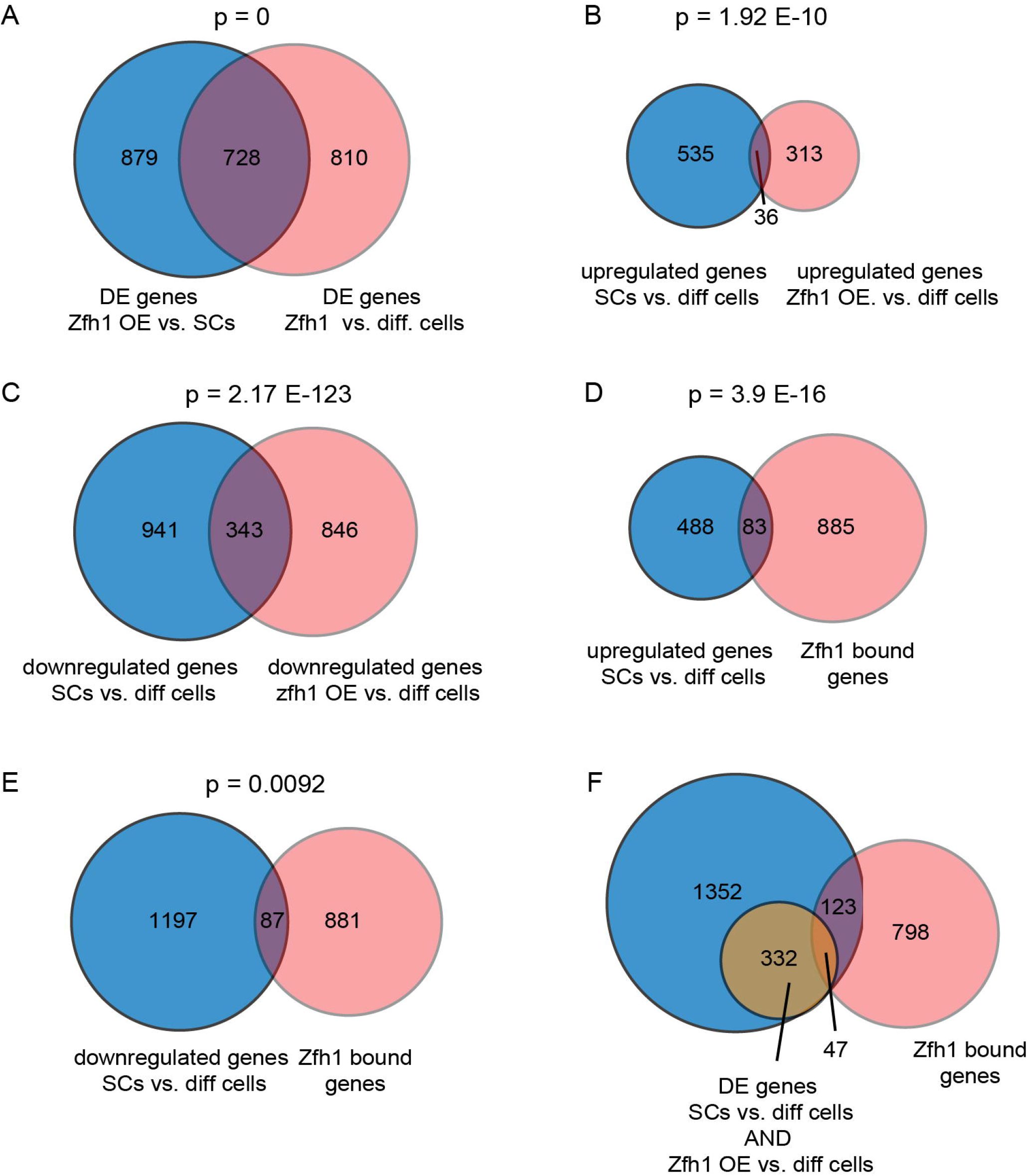
Differential gene expression between somatic cell populations of the testis I. (A-F) Venn diagrams illustrating the overlap between the indicated sets of differentially expressed (DE) genes (A-C) or DE genes and Zfh1 associated genes in the CySCs (D-F). p values, hypergeometric tests.

Nevertheless, the set of 571 genes upregulated in the Zfh1 positive stem cells relative to the differentiated CyCs (suppl. Table S4), shares an overlap of 36 genes with the 349 genes upregulated in tj-positive CyCs overexpressing Zfh1, significantly more than expected (p=1.92 E-10, hypergeometric test) (Fig. 3B). The 1284 and 1245 genes downregulated in stem cells and Zfh1 overexpressing cells, respectively, relative to differentiated CyCs (suppl. Tables S3 and S4) similarly show a larger than expected by chance overlap of 343 genes (p= 2.17 E-123, hypergeometric test) (Fig. 3C), confirming the role of Zfh1 as an important regulator of a subset of cyst stem cell gene activity.

Consistently, 83 of the 571 genes upregulated in somatic stem cells relative to their differentiated progeny and 87 and 1284 of the corresponding, downregulated genes, were also present in the previously identified set of 986 genes bound by Zfh1, in both cases more than expected by chance (p = 3.91 E-16 and p = 0.0092, respectively, hypergeometric test) (Fig. 3 D,E) (suppl. Table S6). Note that these fractions are conservative estimates of the true degree of enrichment, as Zfh1 association was scored solely on basis of the distance between Zfh1 binding site and transcription start (Albert et al., 2018), and thus most likely overestimates the number of Zfh1 regulated genes.

However, only 47 of the 170 Zfh1 associated genes exhibiting differential expression between Zfh1 negative CyCs and endogenously Zfh1 positive CySCs also showed a parallel, consistent response to Zfh1 overexpression, potentially reflecting direct regulation (suppl. Table S6) (Fig. 3F). Genes in this class included e.g. the canonical cyst cell differentiation marker Eya (Fabrizio et al., 2003) (Fig. S5A). For the majority of these genes, however, the changes in transcription levels associated with differentiation appear not to be directly or autonomously set by Zfh1 (Fig. 3F).

### Functional classification of differentially expressed genes in the somatic lineage

We next classified differentially expressed genes in each cell type by gene ontology (GO) analysis using the web based shinyGO package (Ge et al., 2019), based on the full sets of differentially expressed genes as originally called by the DESeq2 algorithm. This revealed 76 biological process GO terms significantly enriched (FDR < 10^-6^) amongst genes overexpressed in differentiated CyCs relative to the CySCs they are derived from (Table S7). Grouping these GO terms into several clusters based on shared gene content reveals a strong overrepresentation of process terms associated with energy metabolism and mitochondrial ATP biosynthesis, in particular including enzymes of the electron transport chain and TCA cycle. Consistently, a separate enriched cluster was comprised of terms related to mitochondrial biosynthesis and protein localization to the mitochondrial matrix (Fig. 4A).

**Figure 4:**
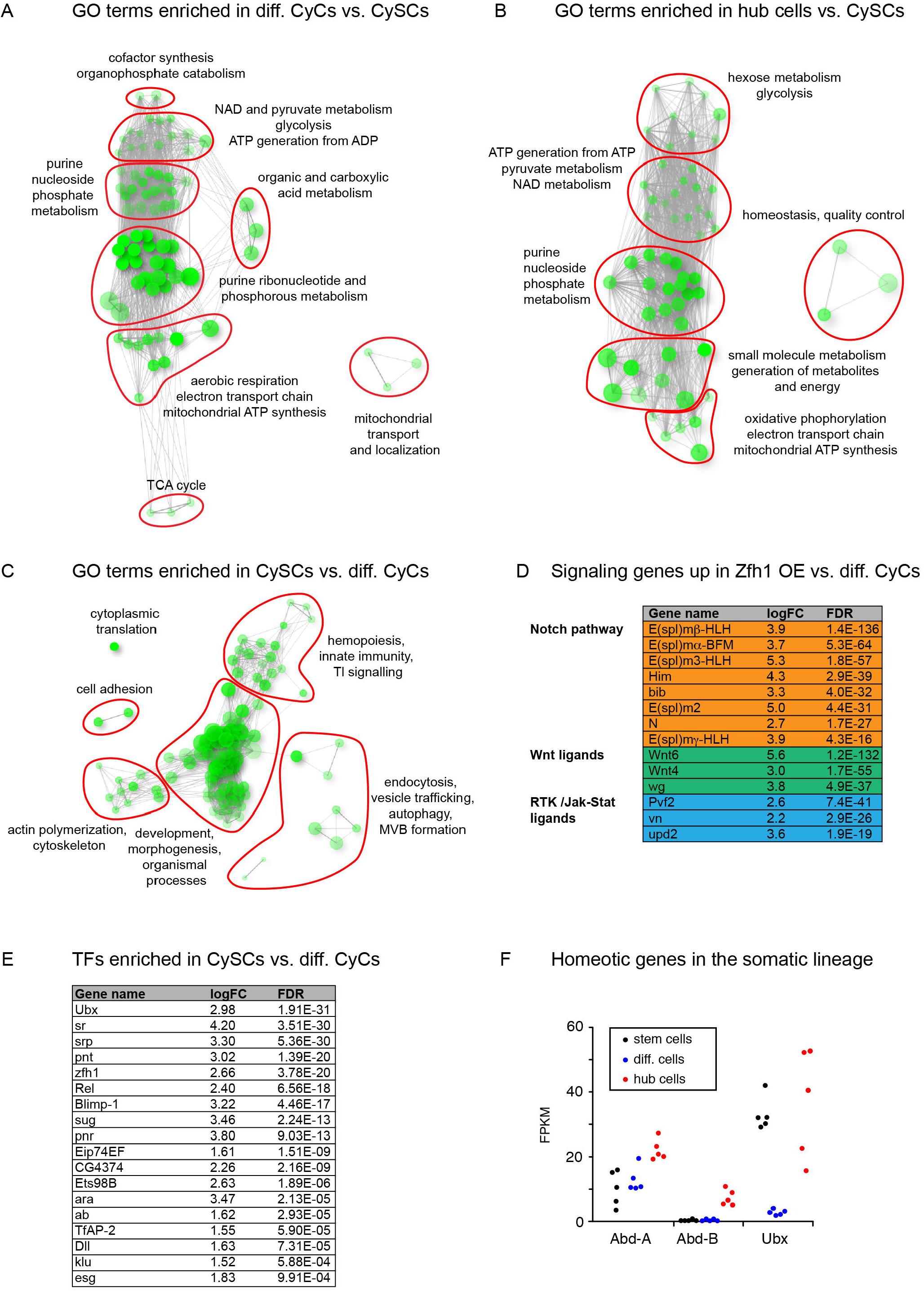
Differential gene expression between somatic cell populations of the testis II. (A-C) Network plots for significantly overrepresented biological process GO terms associated with genes differentially expressed in differentiated CyCs (A), hub cells (B), and CySCs (C). Node size reflects size of the associated gene sets, darker nodes indicate lower FDR, and edge thickness reflects fraction of genes shared between nodes. Red outlines group nodes into clusters with related biological process GO terms. (D) List of signaling genes upregulated in Zfh1 overexpressing cells relative to CyCs. (E) List of transcription factors upregulated (log_2_(fold change) > 1,5) in CySCs relative to differentiated CyCs. (F) Expression levels of homeotic genes in the somatic lineage in FPKM. Each data point corresponds to one replica.

While this may indicate a metabolic specialization of the differentiated CyCs, comparing stem cells and hub revealed a similar pattern, with 58 significantly enriched terms (FDR < 10^−4^) (Table S8) forming clusters associated with hexose catabolism, ATP and pyruvate metabolism, generation of metabolites and energy, as well as oxidative phosphorylation (Fig. 4B). Thus, the set of GO terms related to ATP generation by oxidative phosphorylation may be better described as downregulated in the CySCs relative to the two other somatic cell types.

In contrast, genes differentially expressed in the CySCs were significantly associated with 96 GO terms (FDR < 10^−6^) (Table S9) that formed a large cluster associated with general development and morphogenesis, plus smaller clusters containing terms associated with actin polymerization, endocytosis and other processes involving vesicle trafficking, cell adhesion, and hemopoiesis grouped with innate immune response to pathogens (Fig. 4C). The last of these clusters was rather unexpected, and is therefore the subject of a separate, accompanying manuscript (Hof-Michel and Bökel 2020b, in preparation).

### Zfh1 overexpression induces a unique, tumour like state distinct from both CySCs and differentiated cells

At the level of individual genes, Zfh1 overexpression in the somatic lineage of the testis appeared to induce major changes in intercellular signalling, including e.g. Notch (N) pathway activation and increased production of both Wnt and RTK pathway ligands (Fig. 4D): The top 50 genes by FDR with increased expression in Zfh1 overexpressing cells relative to differentiated CyCs included N itself, the aquaporin-like N pathway member Bib, the E(Spl) complex members Mβ, Mα, M3, M2, and Mγ-HLH, the mesodermal N target Him, as well as the Wnt family ligands Wnt6, Wnt4, and wg, the RTK ligands Pvf2 and vein, and the cytokine-like growth factor Upd2 (suppl. Table S3). This overall picture was mirrored in the comparison with the somatic stem cells (suppl. Table S3), confirming that the Zfh1 overexpressing cells are equally distinct from either endogenous population and therefore of little use as a stem cell proxy. For the remainder, we therefore focused on the genes differentially expressed within the somatic lineage under physiological circumstances.

### Differential expression uncovers additional genes involved in stem cell maintenance

Returning to the strict threshold for differential expression (FDR < 0.001 and |log2(fold change)| > 1.5), the set of 571 genes upregulated in CySCs relative to differentiated cells contained 18 of the 629 annotated PolII transcription factors in the fly genome (Flybase gene group FBgg0000745) (no significant enrichment or depletion, p=0.8284, hypergeometric test) (Fig. 4E). CySCs in the adult testis were distinct from both CyCs and hub cells in their pattern of Hox gene expression: While Abd-A was expressed in all three cell populations, Abd-B remained restricted to the hub, and Ubx expression to hub and stem cells (Fig. 4F).

Most of the transcription factors enriched in the CySCs had not been previously linked to stem cell or niche signalling functions in the somatic lineage of the testis. To test whether our transcriptome analysis had in principle been able to uncover additional transcriptional regulators involved in stem cell function we chose the Drosophila GATA factor Srp (Rehorn et al., 1996) from this list. We generated RFP labelled MARCM clones (Lee and Luo, 1999) homozygous for either a control chromosome or the loss-of function allele *srp*^PZ01549^ (Fig. 5A,B), and assayed whether homozygous clones lacking Srp function were retained in the Zfh1 positive population. Similar to clones lacking the ability to transmit Hh or Upd niche signals (Amoyel et al., 2013; Amoyel et al., 2014; Michel et al., 2012), the fraction of srp^PZ31549^ clones retained in the Zfh1 positive population that contains the stem cells was strongly reduced relative to control by 3d after clone induction (ACI) (Fig. 5C), demonstrating that Srp is indeed required for the maintenance of stemness.

**Figure 5:**
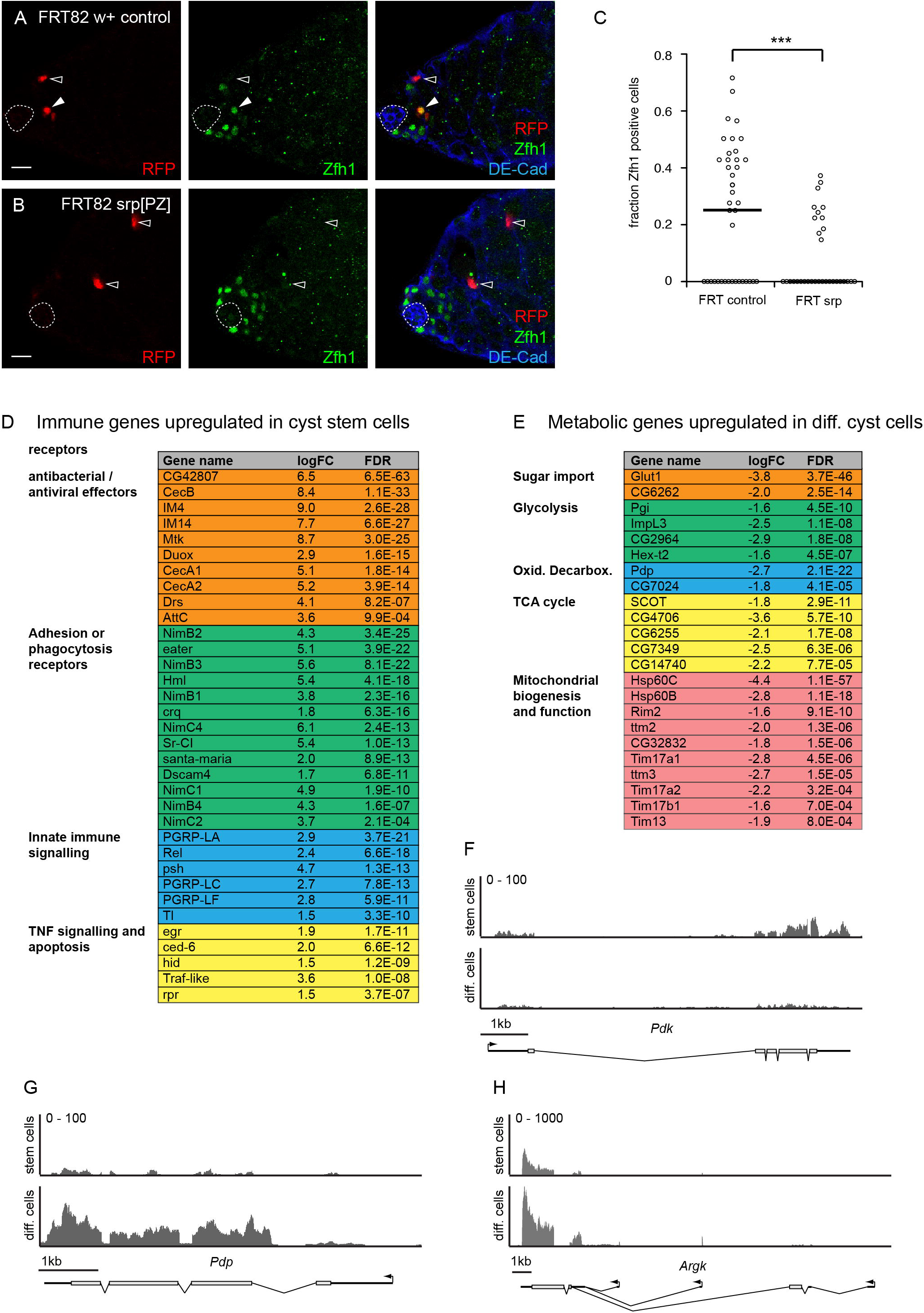
Differential expression and stem cell and cyst cell physiology. (A-C) In contrast to control clones (A), homozygous *srp^PZ^* clones are largely eliminated from the Zfh1 positive (green) stem cell compartment by 3d ACI. Clones marked by RFP (red), cell outlines and hub marked by DE-Cad (blue), hub indicated by dashed outline. Filled and open arrowheads mark Zfh1 positive and negative clonal cells, respectively. (C) Quantification of the Zfh1 positive fraction amongst all clonal cells. Each data point represents one testis. Bar marks median, ***, p<0.001, Mann-Whitney U-test (D) List of genes associated with innate immunity related processes differentially expressed in CySCs (E) List of genes associated with glucose and energy metabolism related processes upregulated in differentiated CyCs (F-H) RNAseq reads plotted against genomic position reveals increased PDK expression in CyCs (F) and increased expression of Pdp (G) and Argk (H) in the differentiated cells. Y axis, read number. (A,B) Scale bars 10μm.

### CySCs but not their differentiated progeny activate an innate immune response during isolation

Genes upregulated in the somatic CySCs relative to their differentiated progeny included 34 genes associated with immune defense, NF-κB or TNF signalling, phagocytosis, or apoptosis, 9 of these within the top 50 hits by FDR, suggesting that the stem cells but not the differentiated CyCs are primed to mount a pathogen response during the isolation procedure (Fig. 5D) (Fig. S5B). However, as reported in detail in the accompanying manuscript (Hof-Michel et al., 2020), this expression pattern does not appear to reflect ongoing, endogenous, antimicrobial activity in the testis. Instead, the differential ability to activate innate immune signalling seems to be related to cell competition (Bowling et al., 2019; Madan et al., 2018) eliminating “loser” stem cells with reduced cellular fitness.

### Differential expression of genes associated with glucose and energy metabolism is relevant for stem cell physiology

As summarized by the GO analysis, the differentiated CyCs showed upregulation relative to the stem cells of multiple biochemical pathways associated with an oxidative phosphorylation biased metabolic state (Fig. 5E). These included mitochondrial transmembrane transporters, proteins involved in mitochondrial biogenesis and function, as well as multiple enzymes of the core glycolysis, oxidative decarboxylation and TCA pathways. In addition to the metabolic enzymes, pyruvate decarboxylase kinase (Pdk), the key negative regulator of pyruvate recruitment by the mitochondrial TCA cycle was downregulated in the differentiated cells (Fig. 5F), while its antagonist pyruvate decarboxylase phosphatase (Pdp) was correspondingly upregulated (Fig. 5G). Importantly, a large number of enzymes of the glucose and energy metabolism missing from the list of differentially expressed genes derived using the strict criteria had been scored as downregulated in stem cells under the initial, relaxed DESeq cutoff (suppl. Table S9). Finally, differentiated CyCs exhibit a threefold increase in arginine kinase (Argk) expression from an already considerable expression level in the stem cells (Fig. 5H). Since arthropods use arginine and Argk as their phosphagen system for buffering ATP levels (Ellington, 2001), this points further at a metabolic differentiation between the two cell types.

To test whether these differences in gene expression also resulted in measurable differences in cellular physiology between CySCs and differentiated CyCs we decided to monitor the oxidative state of the mitochondrial matrix in the two cell types. For this, we first mapped the distribution of mitochondria in the somatic lineage by expressing a GFP fused to a mitochondrial targeting sequence (UAS-mitoGFP) under tj-Gal4 tub-Gal80_ts_ control (Fig. 6A). The granular distribution of the GFP confirmed presence of mature mitochondria already in the CySCs, consistent with prior observations (Senos Demarco and Jones, 2019). We next expressed the UAS-mitotimer sensor (Laker et al., 2014) in the somatic lineage using the same driver line. Mitotimer consists of a mitochondrially targeted RFP featuring a slow transition from the green fluorescent precursor to the mature red fluorescent form. This conversion depends on an oxidation reaction that extends the conjugated pi electron system of the chromophore, and requires the presence of free O_2_ or reactive oxygen species (Verkhusha et al., 2004). Mitotimer was developed to track autophagy mediated mitochondrial turnover through a pulse - chase strategy, with the signal from new born mitochondria dominated by the green precursor fluorescence, while the sensor protein previously localized to older mitochondria already had time to be oxidized to the red fluorescent form (Hernandez et al., 2013). We expressed UAS-mitotimer in a single overnight pulse followed by immediate fixing and staining of the testes. This allowed using the degree of precursor oxidation, as measured by ratiometric imaging of the red and green fluorescence, as a readout for the oxidative state of the mitochondrial matrix in the different cell types (Fig. 6B-D). As predicted from the differential expression of genes associated with mitochondrial ATP generation by oxidative phosphorylation (Fig. 5E), and in particular the key regulators of mitochondrial pyruvate import and oxidative decarboxylation PDK and PDP (Fig. 5F,G), the mitochondrial matrix of differentiated CyCs appeared to provide a more oxidative environment than that of the CySCs. In contrast, mitochondria in the CySCs appeared to be less oxidative (Fig. 6C,D), presumably reflecting a glycolytic rather than oxphos biased metabolic mode.

**Figure 6:**
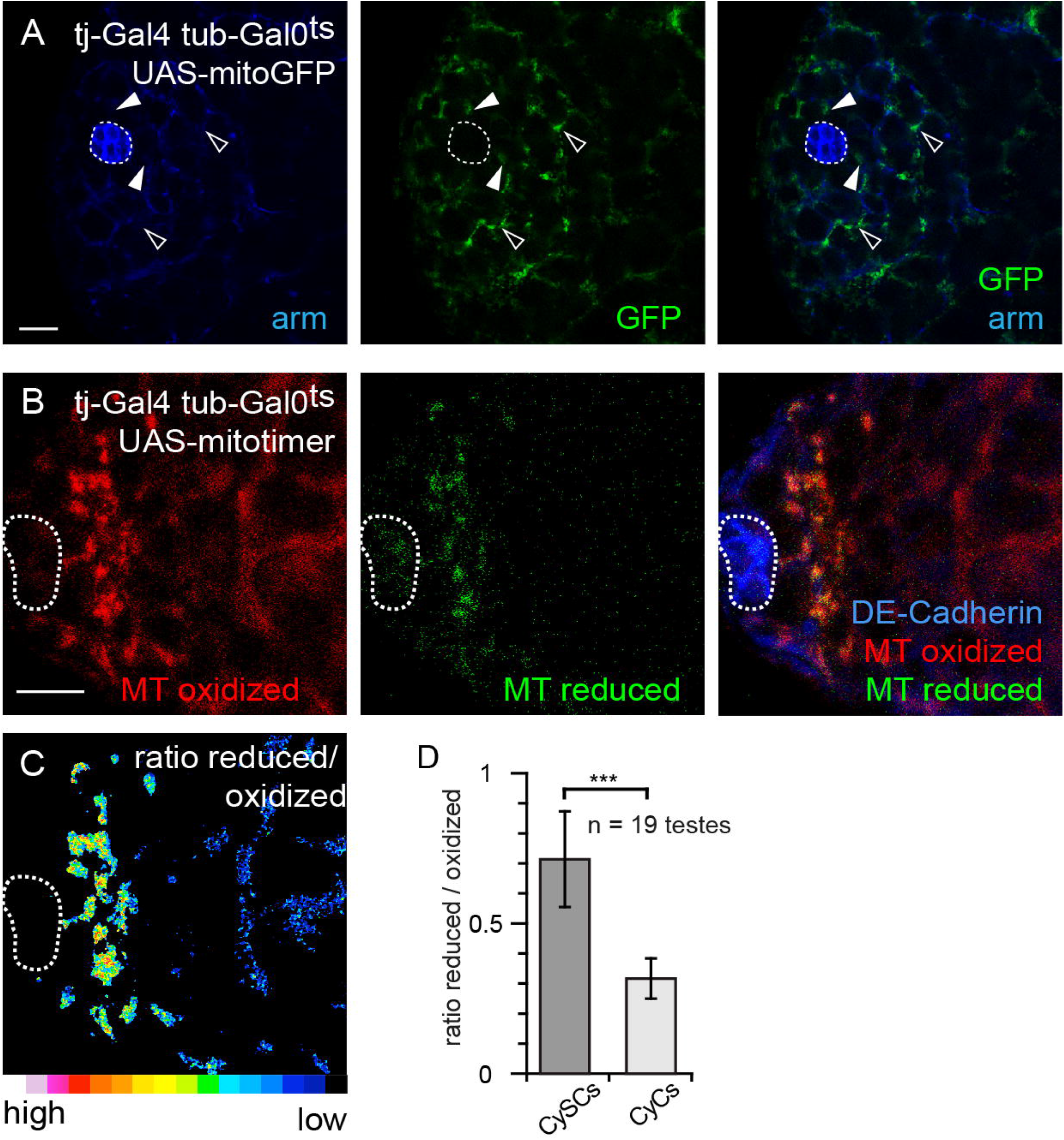
Mitochondrial oxidative state in stem and differentiated cells. (A) Mitochondrially targeted GFP (green) expressed under tj-Gal4 control reveals the distribution of mitochondria in somatic cells of the testis. Hub and cell outlines marked by armadillo (blue), hub marked by dashed outline. Solid arrows, CyCs in the immediate vicinity of the hub, open arrows, differentiated CyCs. (B-D) Measuring the oxidative state of the mitochondrial matrix by pulsed expression of UAS-mitotimer under tj-Gal4 tub-Gal80^ts^ control. (B) Mitotimer is present in a mature oxidized (red) and reduced precursor form (green). Hub marked by DE-Cad (blue) and dashed outline. (C) Ratiometric imaging reveals greater retention of the reduced precursor form in cyst stem cell mitochondria, as quantified in (D). Columns and error bars, mean ± SD, ***, p<0.001, t-test.

The somatic tissues of the testis tip are largely normoxic, as reflected by the effective degradation of a GFP coupled to the O_2_ sensitive ODD domain from Drosophila HIF-α / Sima relative to an RFP control (Misra et al., 2017) (Fig. 7A) and the absence of transcriptional activity from a Sima dependent Gal4 driver containing a fragment of the murine LDH promoter (ldh-Gal4) (Lavista-Llanos et al., 2002) (Fig. S5C). Thus, the CySCs appear to be biased toward aerobic glycolysis, consistent with a Warburg-like (Vander Heiden et al., 2009) stem cell metabolism.

**Figure 7:**
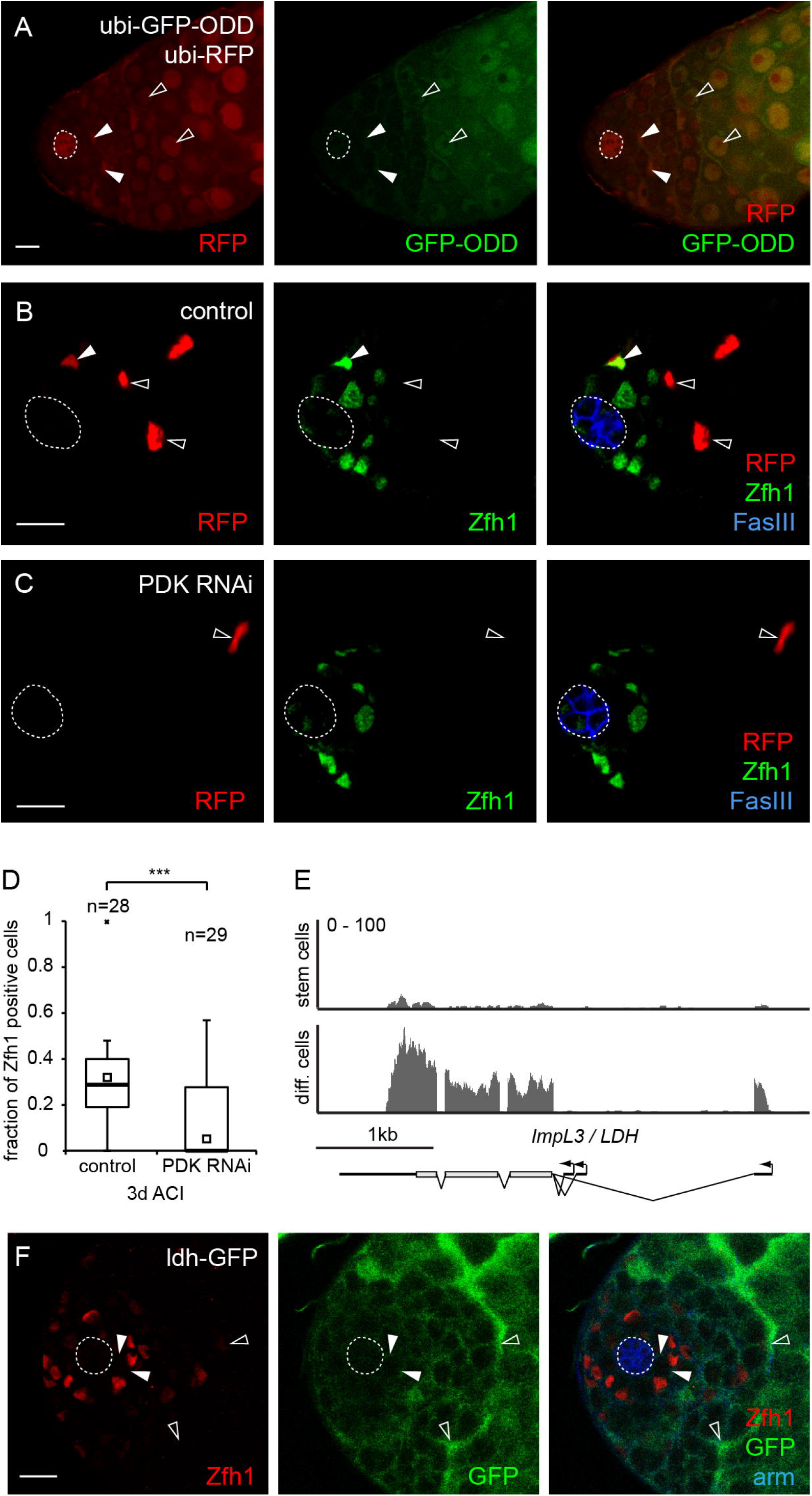
Manipulation of stem cell metabolism leads to stem cell loss. (A) Coexpression of an O_2_-sensitive GFP (green) and RFP (red) under ubiquitin promoter control. GFP degradation demonstrates normoxic conditions in somatic cells near the hub (solid arrowheads). GFP begins to be stabilized further away from the hub (open arrowheads), indicating hypoxic conditions. (B-D) Manipulating stem cell metabolism by clonal PDK knockdown. Control MARCM clones (B) are more efficiently retained in the Zfh1 positive compartment than clones co-expressing a hairpin RNAi construct against PDK (C). (B,C) Hub marked by FasIII (blue), clones marked by RFP (red), Zfh1 (green). Solid arrowheads, Zfh1 pos. clones, open arrowheads, Zfh1 neg. clones. (D) Quantification of the Zfh1 positive cells in (B,C) relative to total clonal cells per testis. (E) Plotting RNAseq reads against genomic position reveals increased ImpL3 / Ldh expression in CyCs relative to CySCs. (F) An ImpL3/Ldh enhancer trap reveals stronger GFP expression (green) in differentiated CyCs (open arrowheads) than in Zfh1 (red) positive cells (solid arrowheads) adjacent to the hub (arm, blue). (A-C,F) Scale bars 10μm, hub marked by dashed outline. (D) Bar, median; small square, mean; box 1st/3rd quartile; whiskers, 1.5x interquartile distance; x, outliers; ***, p<0.001, Kruskal-Wallis test with Tukey’s HSD post hoc test. n, number of testes.

To test this difference for functional relevance we used RNAi knockdown of PDK to clonally force CySCs towards increased pyruvate consumption by the mitochondria (Fig. 7B-D). Compared with controls (Fig. 7B,D), PDK knockdown cells were significantly depleted from the Zfh1 positive population that comprises CySCs and their immediate progeny (Fig. 7C,D), suggesting that a stem cell specific metabolic state is indeed essential for stem cell maintenance.

Finally, even though the above observations suggested that CySCs have a glycolytically biased metabolism when compared with their progeny, the hallmark enzyme of aerobic glycolysis, lactate dehydrogenase (LDH, in *Drosophila* encoded by the *ImpL3* gene) was significantly more strongly expressed in the differentiated CyCs (Fig. 7E). We confirmed this surprising observation using the ldh^YD0852^ GFP enhancer trap line (Quinones-Coello et al., 2007) shown to accurately reflect LDH expression (Wang et al., 2016) (Fig. 7F).

## Discussion

For this, we used genetic labelling of the various cell populations, followed by mechanic and enzymatic dissociation of the tissue and FACS sorting of the cells of interest. While this solved the problem of cell type specificity, the enzymatic step in particular introduced new biases, including e.g. the induction of antibacterial responses due to the experimentally unavoidable exposure of the testis cells to gut contents, that have to be taken into account when interpreting the transcriptome data. Nevertheless, all quality control measures indicated that we had been able to robustly and cleanly isolate the different cell populations, allowing us to draw inferences from the differential expression patterns.

In summary, we have presented, to our knowledge, the first comprehensive and cell type specific transcriptome map of the somatic lineage of the *Drosophila* testis, an established and highly successful model system for stem cell and niche biology (Losick et al., 2011).

The RNAseq method we chose provided a large step up in sensitivity compared with Polll-Dam analysis, the only other method also offering cell type specificity used to date to study transcription in the testis (Tamirisa et al., 2018). In all comparisons, around 10k genes out the ca. 17500 annotated genes in the most recent BDGP genome release (dos Santos et al., 2015) passed the low expression threshold, and between a third to half of these were scored as differentially expressed between the different somatic populations.

We first focused on Zfh1, a transcription factor marking the somatic CySCs and their immediated progeny (Leatherman and Dinardo, 2008). Zfh1 is essential for stem cell maintenance, and has also been considered to be sufficient to induce cyst stem cell state (Leatherman and Dinardo, 2008). However, the Hh niche signal is sufficient to induce Zfh1 expression, but insufficient to establish stem cell state (Amoyel et al., 2013; Michel et al., 2012). Zfh1 instead impinges directly on genetically separable subsets of stem cell behaviours, e.g. limiting proliferation in the somatic lineage to the CySCs by locally repressing members of the Hippo/Warts/Yki pathway (Albert et al., 2018), consistent with models according to which stemness is a compound physiological state rather than a global cell fate determined by some master regulator gene that is turned on or off by the niche (Clevers and Watt, 2018; Post and Clevers, 2019).

Our transcriptome data further supports this notion. Comparing the expression differences between Zfh1-negative, differentiated CyCs and both endogenously Zfh1-positive CySCs and artificially Zfh1 overexpressing CyCs demonstrated that differentially expressed genes are enriched for genes associated with Zfh1 binding sites as measured by *in vivo* Zfh1-DamID in the stem cells (Albert et al., 2018). Nevertheless, only a minority of these genes, including e.g. the cyst cell differentiation marker Eya (Fabrizio et al., 2003), showed consistent, parallel expression changes in response to the endogenous or artificial presence of Zfh1, suggesting that for the majority of target genes Zfh1 by itself is insufficient to set the expression level.

Rather than mimicking a stem cell transcription profile, Zfh1 overexpression instead induced differential expression of thousands of genes, prominently including signatures of altered N and Wnt pathway activity. Indeed, CySCs differ from differentiated CyCs to a similar degree as the Zfh1 induced proliferating cells differ from either endogenous population. The small, Zfh1-positive and Eya-negative tumour-like cells induced by Zfh1 overexpression are therefore of extremely limited use as a stem cell model.

Concentrating instead on the endogenous cell populations, the transcriptome of the CySCs shows a strong signature of antimicrobial defense that is completely missing from their differentiated cyst cell progeny (Fig. S5B). First, this sharp difference confirms that we had cleanly separated differentiated cells from the cyst stem cell sample. Second, since these populations are in direct contact within the testis, it appears implausible that endogenously one but not the other cell type needs to engage in antibacterial defence. In a separate manuscript we therefore argue in detail that activation of the antimicrobial response is an artefact caused by the mechanic and enzymatic disruption of the testis tissue, as no such response is detectable in intact testes. However, the differential ability of the stem cells to activate innate immune signalling is real, and is instead related to competition between stem cells resulting in elimination of the less fit loser cells from the niche (Hof-Michel and Bökel 2020b, in preparation).

GO analysis revealed additional clusters of biological function terms enriched within the genes upregulated in stem cells relative to differentiated cells. These include terms associated with autophagy, vesicle trafficking and endocytosis, recalling observations by multiple labs linking testis stem cell and niche function to genes involved in protein trafficking and endocytosis (Cook et al., 2017; Fairchild et al., 2017; Papagiannouli et al., 2019; Tang et al., 2017). Similarly, another cluster containing GO terms associated with actin and cytoskeletal function is not unexpected due to the established link between niche signalling and stem cell adhesion to the hub mediated by the Rap-GEF *Gef26 / dizzy* (Wang et al., 2006).

Compared to the CySCs, both hub cells and differentiated CyCs were enriched for GO clusters associated with an oxidative phosphorylation based metabolism. Conversely, the relative downregulation of the associated genes in stem cells implies a relative bias towards glycolytic energy production, even though the HIF and oxygen sensor experiments suggested that the stem cell region at the testis tip is largely normoxic. This pattern would be characteristic of a Warburg-type, aerobic glycolysis biased stem cell metabolism (Vander Heiden et al., 2009), which is also supported by the less oxidative state of the stem cell mitochondria implied by our Mitotimer sensor data. This metabolic mode seems to be essential for stem cell maintenance, as forcing pyruvate uptake by the mitochondria by PDK knockdown is sufficient to drive their elimination form the stem cell pool. However, it is possible that this metabolic challenge renders these cells “unfit” relative to their unaffected neighbours, thus inducing their elimination as “losers” in the ongoing stem cell competition, even though they may, in principle, still be able to function as stem cells. In the future, these questions will have to be revisited, using additional sensors to directly monitor the metabolic changes in the affected cells and employing genetic tools to abolish competition, which would exceed the scope of this manuscript.

Interestingly, though, most glycolytic enzymes, including the hallmark gene ImpL3 encoding lactate dehydrogenase, are consistently also upregulated in the differentiated cells. It is therefore tempting to speculate that differentiated CyCs generate their own energy by oxidative phosphorylation, as marked by the upregulation of genes associated with mitochondrial biogenesis and function and the relatively more oxidative state of their mitochondria. In addition they may, as part of their blood-germline barrier duties (Fairchild et al., 2015; Fairchild et al., 2016) use Ldh to provide energy carriers to the developing germline clusters they ensheath, similar to the metabolic division of labour between neurons and glia (Volkenhoff et al., 2015). Again, these ideas will in the future have to be tested using additional sensors and genetic manipulations that would exceed the scope of this manuscript.

We hope, however, that the present transcriptome data will already be a useful resource for the fly stem cell community, stimulating further research using this successful and experimental tractable model system.

## Materials and Methods: Fly stocks and transgenic constructs

w; P{w[+mC]=arm-lacZ.V}36BC P{ry[+t7.2]=neoFRT}40A/CyO, P{ry[+t7.2]=sevRas1.V12}FK1 (BL-7371), w; P{ry[+t7.2]=neoFRT}42D P{w[+mC]=arm-lacZ.V}51D (BL-7372), w; P{w[+mC]=UAS-RedStinger} (BL-8547), w; P{ry[+t7.2]=neoFRT}82B P{w[+t*] ry[+t*]=white-un1}90E (BL-2050), and w; P{w[+mC]=UAS-MitoTimer} (BL-57323) were obtained from the Bloomington Drosophila Stock Center and have been described. The w hs-FLP C587-Gal4 UAS-RedStinger chromosome has been described before (Michel et al., 2012). The following stocks were graciously provided by our colleagues: P{ry[+t7.2]=PZ} srp^01549^/ TM3,ry Sb Ser (srp^PZ^, Ingolf Reim, FAU Erlangen), Pin/CyO;Gal4-Gal80Hack/TM6B (94E5) (Christopher Potter, Johns Hopkins Univ., Baltimore), w;; ldh-GFP[YD0852] (Utpal Banerjee, UCLA), w; UAS-RFP UAS-ODD-GFP and w; ldh-Gal4 (Stefan Luschnig, Univ. Münster).

w;; zfh1-T2A-T2A-Gal80 (referred to in brief as zfh1-Gal80) was generated by crossing yw vasa-Cas9 first to zfh1-T2A-Gal4 w+ / TM3, Sb (Albert et al., 2018) and then to the Pin / CyO; Gal4-Gal80Hack / TM6B (94E5) Gal4-Gal80 HACK stock (Lin and Potter, 2016) that contains all the required components such as gRNA genes, homology arms, and an eye RFP selection marker to insert a T2A-Gal80 cassette into the Gal4 ORF of any Gal4 transgene on the homologous chromosome.

MARCM clones (Lee and Luo, 1999) were generated by crossing FRT males, where required also carrying additional UAS or reporter constructs, to w hs-FLP C587-Gal4 UAS-RedStinger virgins carrying the matching tub-Gal80 FRT chromosome and heat shocking adult males for 30min h at 37°C. Clones were generated using the following FRT chromosomes: arm-lacZ FRT40A, FRT42D arm-lacZ, FRT82 w^+^90E, and FRT82 srp^PZ01549^.

The following stocks or crosses were used to label somatic cell populations in the testis: upd-Gal4 tub-Gal80^ts^;;UAS-RedStinger (hub cells), w;;zfh1-T2A-Gal4/TM3,Sb x w;;UAS-RedStinger (CySCs), tj-Gal4 tub-Gal80^ts^/CyO; UAS-RedStinger/MKRS x w;;zfh1-T2A-T2A-Gal80w+/MKRS (differentiated CyCs), and tj-Gal4 tub-Gal80^ts^/CyO; UAS-RedStinger/MKRS x w;;UAS-zfh1 (Zfh1 induced tumours).

### Antibodies and immunohistochemistry

Testes were stained as described (Albert 2018). The following antisera were used: rat-anti-DECadherin (DSHB, DCAD2) 1:100, mouse-anti-FasIII (DSHB, 7G10) 1:250, mouse-anti-Armadillo (DSHB, N2 7A1) 1:100, rabbit-anti-Zfh1 (Ruth Lehmann) 1:4000. Goat secondary antisera labelled with Alexa-488, −568, or −633 (Invitrogen) were used 1:500.

### Imaging and image analysis

Images were acquired using a Leica SP5 confocal microscope with a HCX PL APO 40x / NA1.25 oil immersion objective. Unless stated otherwise, images are single optical slices. Image quantifications were performed using Fiji (Schindelin et al., 2012). Differences between samples were tested for significance as appropriate with Student’s t-test, Mann-Whitney U-test or Kruskal-Wallis test followed by Tukey’s HSD, using the PMCMRplus R package (Pohlert, 2018). Images were prepared for publication using Adobe Photoshop and Illustrator.

### Cell isolation and purification

Flies with cell type specific expression of nuclear RFP in the somatic lineage of the testis (hub cells, somatic stem cells, differentiated CyCs, or Zfh1 overexpressing tumour cells) were hand dissected on ice in Schneider’s medium with L-Glutamine (Pan Biotech), separating the testes proper from the remainder of the gonad (accessory glands, ejaculatory duct, etc.). 20-30 testes per sample were transferred to a 2ml cryotube (Nunc). Following aspiration of the medium, testes were covered with 1.5 ml separation buffer (Schneider’s medium with L-Glutamine, 8.25μg/μl Collagenase, 0.005 μg/μl Trypsin, 1.75 mμ EDTA) and vigorously agitated on a shaker at 30°C for 30 min. The medium was then replaced with ice cold PBS, and individual cells released from the debris by repeated pipetting (100x) using 20-300μl tips (epT.I.P.S. LoRetention Dualfilter, Eppendorf) followed by filtering through a 40 μm cell strainer (Falcon).

Finally, 150 cells per replica were sorted directly into a 384 well skirted plate (Eppendorf twintec 0030128648) containing per well 2 μl lysis buffer comprised of 1.9 μl of 0.2% Triton X-100 diluted in nuclease free H_2_O (Invitrogen 10977049) and 0.1 μl 40 U/μl murine RNase Inhibitor (NEB M0314S), using a Beckman Coulter MoFlo Astrios sorter at the Flow Cytometry Core Facility at Philipps-University Marburg. Following sorting, cells were spun down and immediately frozen on dry ice.

### RNAseq and bioinformatics

PolyA(+) mRNA was isolated from the samples, reverse transcribed, and amplified (15-18 cycles) by the Deep Sequencing Core Facility of the Center for Molecular and Cellular Biolengineering at the Technical University Dresden using the SmartSeq2 (Illumina) kit and protocol (Picelli et al., 2014). Libraries were then generated using the Nextera kit (Illumina) and sequenced to a raw target depth of 10Mio 75bp single end reads using an Illumina NextSeq500 sequencer.

Reads were quality checked and mapped to the *Drosophila melanogaster* dm6 reference genome (dos Santos et al., 2015) with the help of the A.I.R. RNAseq web based analysis package (Sequentia Biotech, Barcelona). The underlying algorithms and software packages collated in A.I.R. are described in (Vara et al., 2019).

The low expression cutoff for inclusion of genes into pairwise transcriptome comparisons, as implemented in the A.I.R. package, was set by calculating a global Jaccard index for each sample type, progressively eliminating genes with the smallest FPKM values until correlation between replicates was maximized. To assign differential gene expression between samples we chose the DESeq algorithm (Love et al., 2014) as implemented in the A.I.R. package with an default FDR cutoff of 0.05.

Additional bioinformatic analyses (heatmaps, Spearman correlation plots, Volcano plots, Venn diagrams and associated hypergeometric tests) were performed using RStudio, relying in particular on the bioconductor R package (Huber et al., 2015).

Enrichment of gene ontology terms for the various differentially expressed gene sets was assayed using the shinyGO web platform (Ge et al., 2019).

## Supporting information

Table S1 Mapping statistics

Table S6 DE genes wirh consistent Zfh1 response

Table S2 expression levels all genes

Table S3 DE genes in Zfh1 OE

Table S4 DE genes CySCs vs. diff. CyCs

Table S5 DE genes CySCs vs. hub

Table S7 GO terms up in diff CyCs vs. CySCs

Table S8 GO terms up in hub cells vs. CySCs

Table S9 GO terms up in CySCs vs. diff CyCs

Table S10 DE metabolic genes

## Data availability

Processed RNA-seq results are presented in the supplementary information. The corresponding raw data will be available after publication at the NCBI Sequence Read Archive (BioProject xxxxx).

## Author contributions

SHM established the cell purification protocol and performed initial fly experiments, imaging, and analyses of imaging data. CB performed molecular cloning, fly and imaging experiments, was responsible for imaging and RNAseq data analysis, devised the project, and wrote the manuscript.

## Acknowledgements

We would like to thank Christopher Potter, Ingolf Reim, Ruth Lehmann, Stefan Luschnig, Utpal Banerjee, and the JEDI community for fly stocks, reagents, and advice, Erika Bach for critical comments on the manuscript, Ljubinka Cigoja (fly genetics and immunocytochemistry) and Sabina Huhn (fly genetics and molecular cloning) for their expert technical assistance, Isolde Kranz for supplying fly food and coffee, Eugene Albert for his initial work on the cell isolation protocol, Gavin Giel and the Flow Cytometry Core Facility at Philipps-University Marburg for sorting our cells, and Andreas Dahl and Susanne Reinhardt at the CRTD Deep Sequencing Core Facility for the RNA sequencing. The project was supported by DFG grant BO 3270/3-1 to CB.

