## Supplementary Figures S1-S4 for "Transcriptome analysis of somatic cell populations in the *Drosophila* testis links metabolism and stemness"

**Supplementary Information Hof-Michel and Bökel, 2020:**

**Supplementary tables S1-S9**

**Table S1:** RNAseq mapping statistics

**Table S2:** Genome wide expression levels in the somatic lineage of the testis

**Table S3:** Genes with differential expression following Zfh1 overexpression

**Table S4:** Genes with differential expression between CySCs and differentiated CyCs

**Table S5:** Genes with differential expression between CySCs and hub

**Table S6:** Differentially expressed genes with consistent Zfh1 response

**Table S7:** GO terms enriched in genes differentially expressed in diff. CyCs vs. CySCs

**Table S8:** GO terms enriched in genes differentially expressed in hub cells vs. CySCs

**Table S9:** GO terms enriched in genes differentially expressed in CySCs vs. diff. CyCs

**Table S10:** Differentially expressed genes associated with glucose and energy metabolism

**Supplementary Figures S1 to S5**

**Figure S1: Zfh1 constructs and isoform specific deletions.** (A) Generating zfh1-T2A-T2A-Gal4 from the zfh1-T2A-Gal4 Crispr/Cas9 knockin line. Zfh1-T2A-Gal4 was made by integrating a $w^+$ marked T2A-Gal4 cassette into the endogenous Zfh1 locus, producing a C-terminal, co-translationally separating fusion protein. Zfh1-T2A-T2A-Gal4 was then generated by introducing an in-frame insertion of T2A-RFP cassette tagged with eye-RFP into the GFP ORF using the Potter lab HACK approach. (B) Isoform specific deletions of Zfh1-RA and Zfh1-RB. Both deletions were generated by Crispr/Cas9. *zfh1*$^{\Delta RA}$ was recovered by injecting two gRNAs flanking the start codon into y w vasa-Cas9 and testing candidate chromosomes for lethality over Df(3R)exel9020 that deletes the entire Zfh1 locus before PCR screening. *zfh1*$^{\Delta RB}$ is viable and was recovered by PCR screening progeny from crossing a dual gRNA transgenic chromosome to the Cas9 source. (C) *zfh1*$^{\Delta RB}$ is viable and fertile over Df(3R)exel9020. Testes of such males are morphologically normal, and Zfh1 immunostaining (red) is indistiguishable from WT due to the remaining isoform Zfh1-RA. Hub labelled with DE-Cadherin (blue), scale bar 10 µm.

**Figure S2: FACS isolation of somatic cell populations in the Drosophila testis.** (A-D) Log-log scatter plots of RFP labelled cells at final sorting. A $w^-$ negative control (A) lacks RFP high cells in the lower right quadrant that appear in the same gate in testes from flies expressing RFP in differentiated cyst cells under tj-Gal4 tub-Gal80ts Zfh1-T2A-T2A-Gal80 control (B). Cells from flies expressing RFP under zfh1-T2A-Gal4 Gal80ts control (C) show distinct RFP low and RFP high populations. Co-overexpression of Zfh1 with RFP under tj-Gal4 tub-Gal80ts control (D) leads to the expansion of a population with very high RFP levels. Gate R3 was used for sorting in all cases (rescaled in D).

**Figure S3: RNAseq validation I.** Spearman correlation between the full transcriptomes of replicates for the four sample types. Note the overall better correlation for samples with higher numbers of labelled cells per testis in the starting material (differentiated and Zfh1 overexpressing cells), even though identical numbers of cells were used for all 20 replicas.

**Figure S4: RNAseq validation II.** Heatmap clustering based Euclidean distance mapping of $\log_2$ fold enrichment scores of genes marked as differentially expressed using the DESeq2 algorithm with the default settings of the Sequentia A.I.R. package (FDR < 0.05). Light blue line, histogram of gene count vs. log2 fold enrichment. For comparisons of CySCs vs. differentiated CyCs (A), Zfh1 overexpressing cells vs. diff. CyCs (B), Zfh1 overexpressing cells vs. CySCs (C), and hub cells vs. CySCs consistent cluster structures are recovered.

**Figure S5: Differentially gene expression and testis HIF activation.** (A,B) Plotting RNAseq reads (A,B) and Zfh1-Dam and Dam only control reads (A) against genomic position reveals increased eya expression in cyst cells relative to cyst stem cells as well as strong Zfh1 binding in stem cells near the eya internal promoter (A), while expression of the antimicrobial peptide Drs is strictly limited to cyst stem cells (B). (C) Nuclear RFP expression (red) under control of a Sima/HIF-α dependent Gal4 driver containing a fragment of the murine LDH promoter (ldh-Gal4) can be detected in rare individual cells of the muscle sheath, but is absent from the vicinity of the hub (marked with FasIII, blue, and a dashed outline) and the Zfh1 (green) positive compartment. (A,B) Y axis reads in FPKM, (C) Scale bar 10 μm.

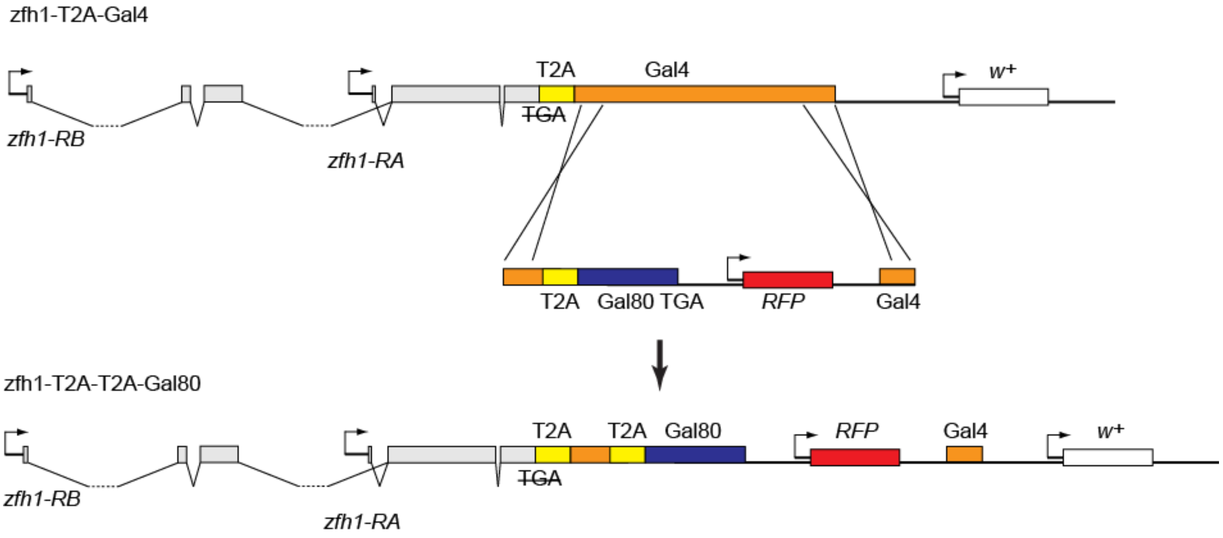

zfh1-T2A-Gal4
T2A
Gal4
w+
TGA
zfh1-RB
zfh1-RA
T2A
Gal80 TGA
RFP
Gal4
zfh1-T2A-T2A-Gal80
T2A
T2A
Gal80
RFP
Gal4
w+
TGA
zfh1-RB
zfh1-RA


**Fig. S1**

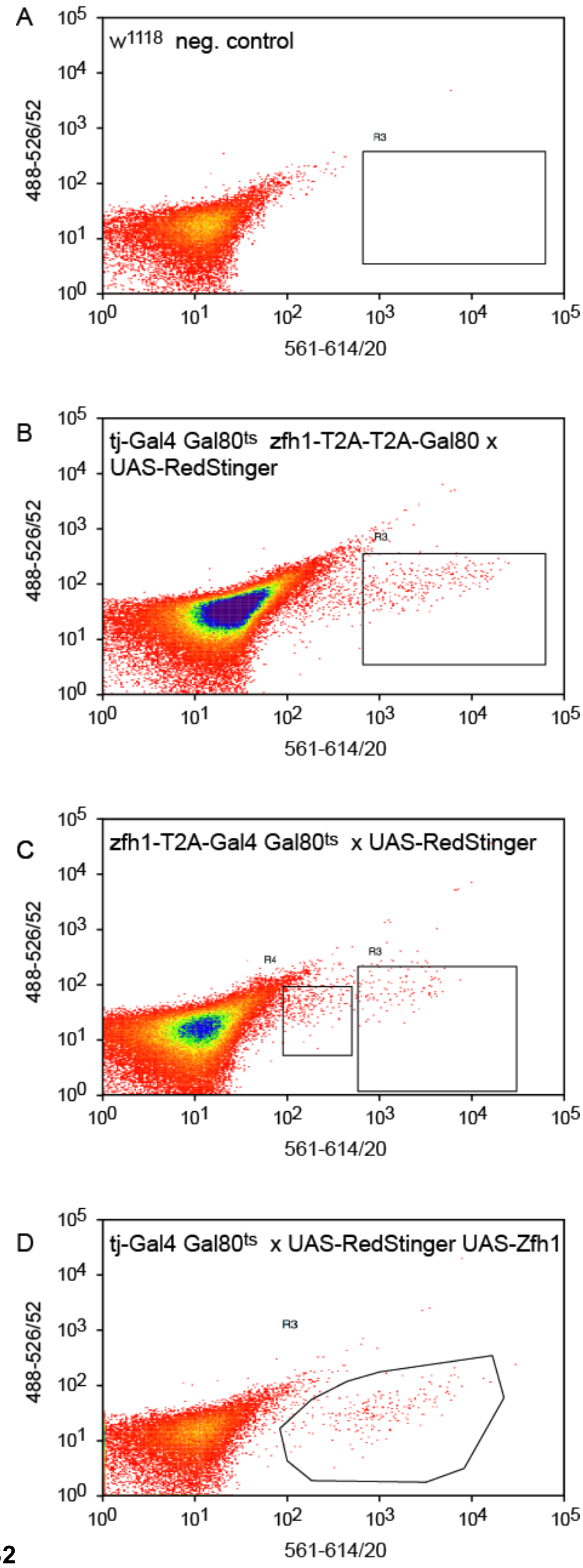
A
w1118 neg. control
R3
B
tj-Gal4 Gal80ts zfh1-T2A-T2A-Gal80 x UAS-RedStinger
C
zfh1-T2A-Gal4 Gal80ts x UAS-RedStinger
R4
D
tj-Gal4 Gal80ts x UAS-RedStinger UAS-Zfh1
488-526/52
561-614/20
10^0
10^1
10^2
10^3
10^4
10^5

**Fig. S2**

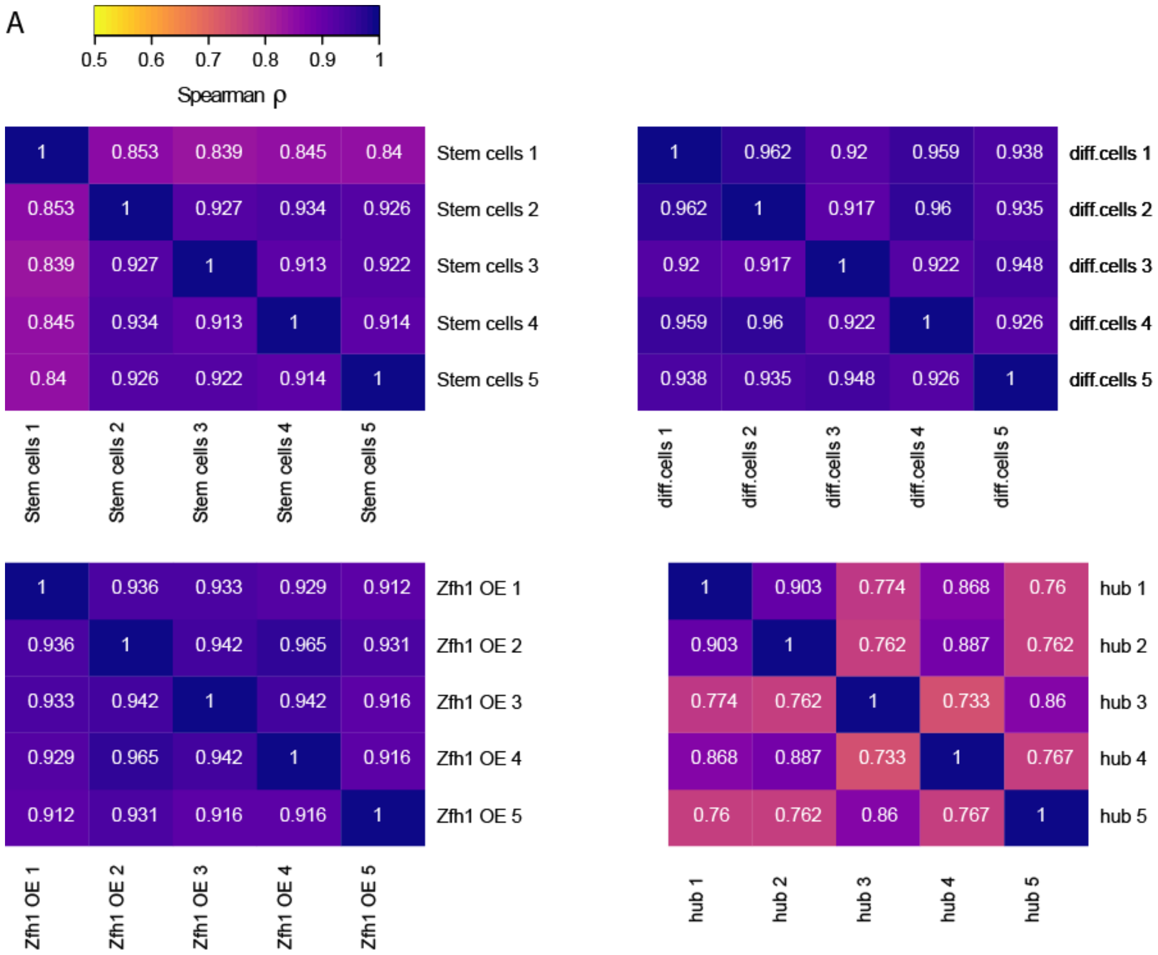
A
0.5 0.6 0.7 0.8 0.9 1
Spearman ρ
1 0.853 0.839 0.845 0.84 Stem cells 1
0.853 1 0.927 0.934 0.926 Stem cells 2
0.839 0.927 1 0.913 0.922 Stem cells 3
0.845 0.934 0.913 1 0.914 Stem cells 4
0.84 0.926 0.922 0.914 1 Stem cells 5
Stem cells 1 Stem cells 2 Stem cells 3 Stem cells 4 Stem cells 5
1 0.962 0.92 0.959 0.938 diff.cells 1
0.962 1 0.917 0.96 0.935 diff.cells 2
0.92 0.917 1 0.922 0.948 diff.cells 3
0.959 0.96 0.922 1 0.926 diff.cells 4
0.938 0.935 0.948 0.926 1 diff.cells 5
diff.cells 1 diff.cells 2 diff.cells 3 diff.cells 4 diff.cells 5
1 0.936 0.933 0.929 0.912 Zfh1 OE 1
0.936 1 0.942 0.965 0.931 Zfh1 OE 2
0.933 0.942 1 0.942 0.916 Zfh1 OE 3
0.929 0.965 0.942 1 0.916 Zfh1 OE 4
0.912 0.931 0.916 0.916 1 Zfh1 OE 5
Zfh1 OE 1 Zfh1 OE 2 Zfh1 OE 3 Zfh1 OE 4 Zfh1 OE 5
1 0.903 0.774 0.868 0.76 hub 1
0.903 1 0.762 0.887 0.762 hub 2
0.774 0.762 1 0.733 0.86 hub 3
0.868 0.887 0.733 1 0.767 hub 4
0.76 0.762 0.86 0.767 1 hub 5
hub 1 hub 2 hub 3 hub 4 hub 5

**Fig. S3**

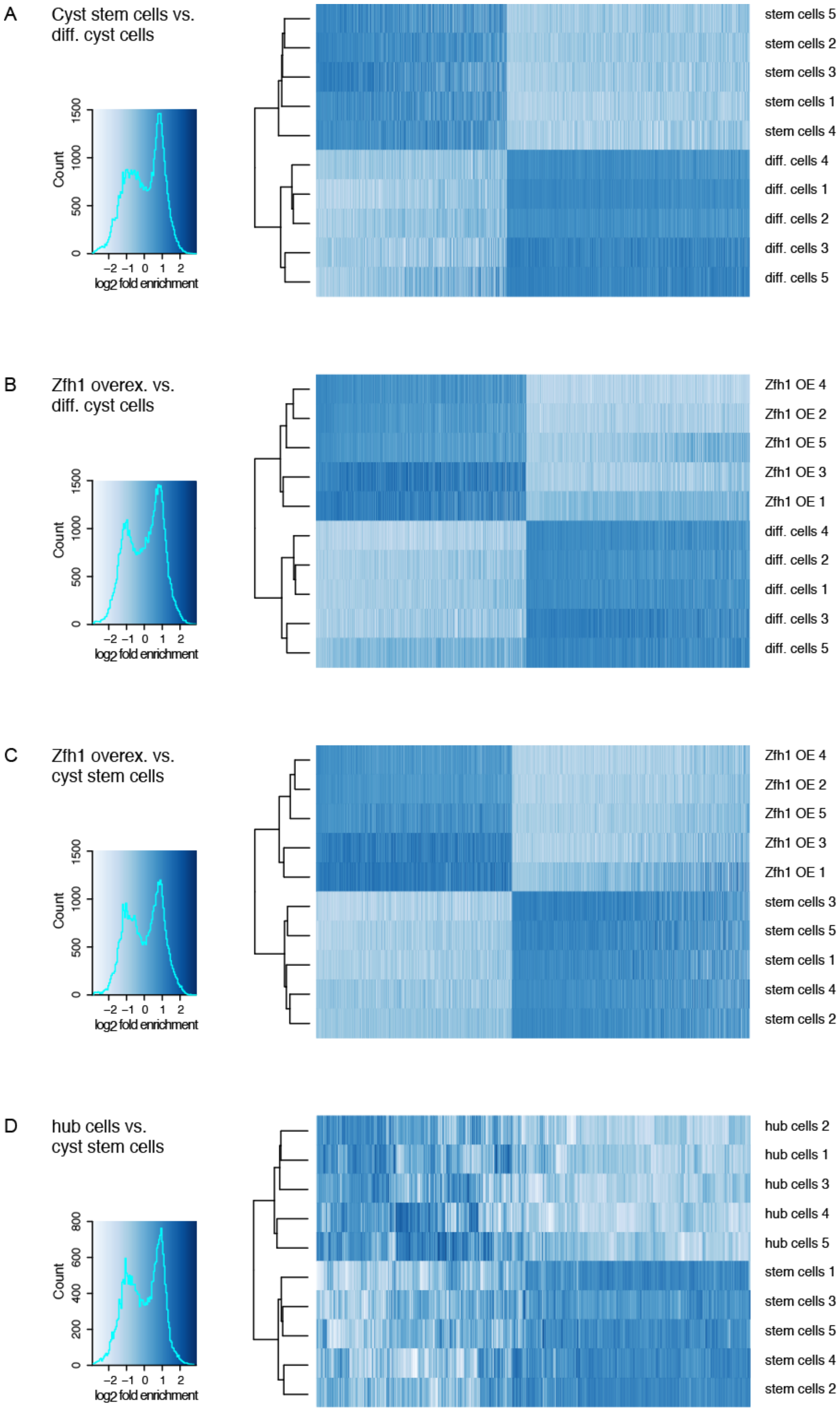
A
Cyst stem cells vs.
diff. cyst cells
Count
log2 fold enrichment
stem cells 5
stem cells 2
stem cells 3
stem cells 1
stem cells 4
diff. cells 4
diff. cells 1
diff. cells 2
diff. cells 3
diff. cells 5
B
Zfh1 overex. vs.
diff. cyst cells
Count
log2 fold enrichment
Zfh1 OE 4
Zfh1 OE 2
Zfh1 OE 5
Zfh1 OE 3
Zfh1 OE 1
diff. cells 4
diff. cells 2
diff. cells 1
diff. cells 3
diff. cells 5
C
Zfh1 overex. vs.
cyst stem cells
Count
log2 fold enrichment
Zfh1 OE 4
Zfh1 OE 2
Zfh1 OE 5
Zfh1 OE 3
Zfh1 OE 1
stem cells 3
stem cells 5
stem cells 1
stem cells 4
stem cells 2
D
hub cells vs.
cyst stem cells
Count
log2 fold enrichment
hub cells 2
hub cells 1
hub cells 3
hub cells 4
hub cells 5
stem cells 1
stem cells 3
stem cells 5
stem cells 4
stem cells 2

**Fig. S4**

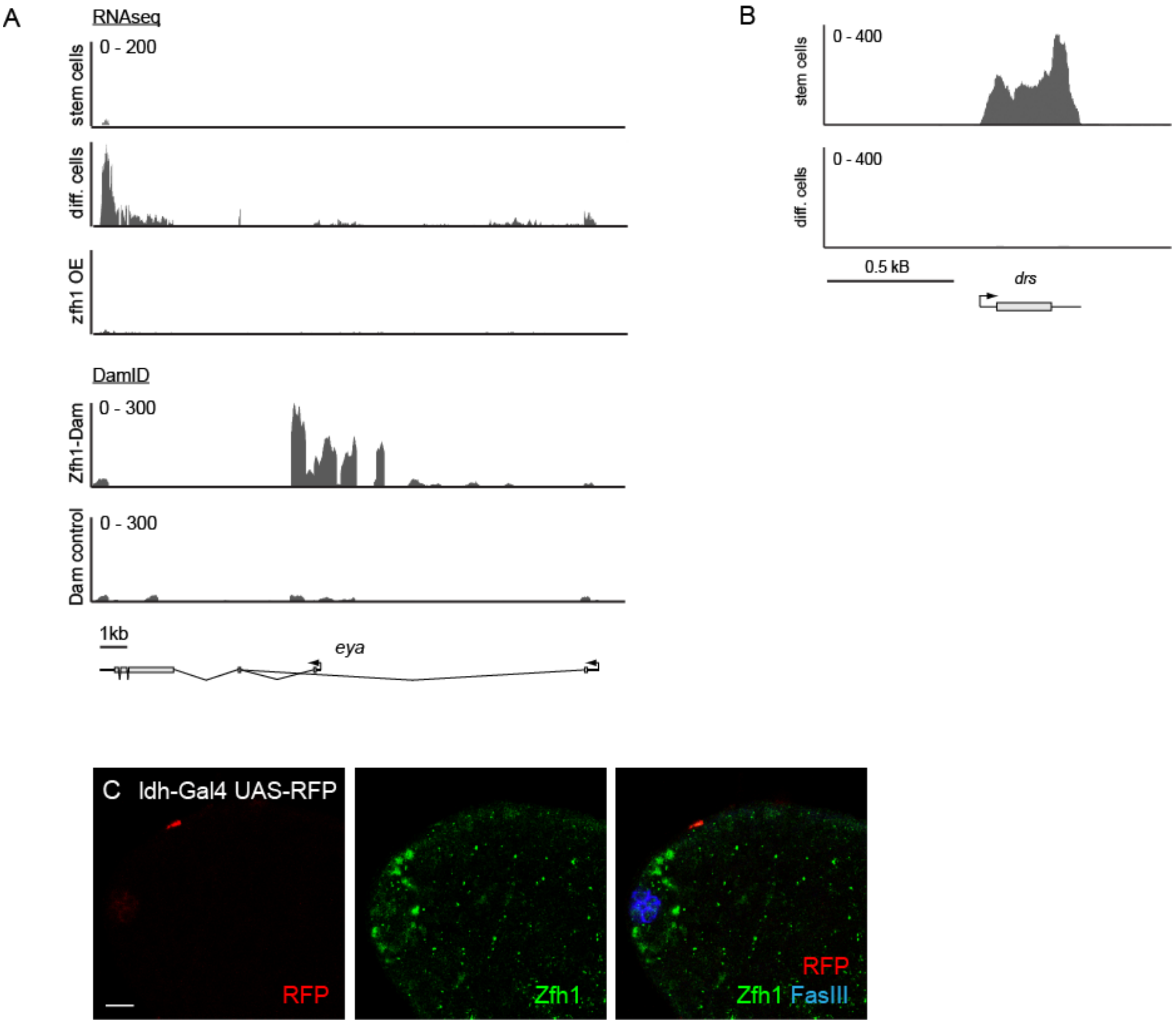
A
RNAseq
0 - 200
stem cells
diff. cells
zfh1 OE
DamID
0 - 300
Zfh1-Dam
0 - 300
Dam control
1kb
eya
B
0 - 400
stem cells
0 - 400
diff. cells
0.5 kB
drs
C Idh-Gal4 UAS-RFP
RFP
Zfh1
RFP
Zfh1 FasIII

**Fig. S5**
